# Wnt and BMP signalling direct anterior/posterior differentiation in aggregates of mouse embryonic stem cells

**DOI:** 10.1101/2023.04.07.535990

**Authors:** Atoosa Amel, Simoné Rossouw, Mubeen Goolam

## Abstract

Stem cell-based embryo models have allowed greater insight into peri-implantation mammalian developmental events that are otherwise difficult to manipulate due to the inaccessibility of the early embryo. The rapid development of this field has resulted in the precise roles of frequently used supplements such as N2, B27 and Chiron in driving stem cell lineage commitment not being clearly defined. Here, we investigate the effects of these supplements on embryoid bodies to better understand their roles in stem cell differentiation. We show that Wnt signalling has a posteriorising effect on stem cell aggregates and directs differentiation towards the mesoderm, as confirmed through the upregulation of posterior and mesodermal markers. N2 and B27 can mitigate these effects and up-regulate the expression of anterior markers. To control the Wnt gradient and the subsequent anterior vs. posterior fate, we make use of a BMP4 signalling centre and show that aggregates in these conditions express cephalic markers. These findings indicate that there is an intricate balance between various culture supplements and their ability to set up the anterior/posterior axis in stem cell embryo models.

**Summary Statement:** The complex reagents used in ’stembryo’ protocols have unclear roles in stem cell differentiation in vitro requiring further investigation. This study examines their effects on embryoid bodies.

## Introduction

Successful mammalian development is dependent on precise communication and coordination between embryonic and extraembryonic tissues, leading a small collection of cells to form a highly complex, precisely defined multicellular organism. The study of these intricate morphogenic events is complicated by the fact that the most critical organising events occur during implantation, hidden from view. Implantation stage development is accompanied by critical cell proliferation, migration, and differentiation events that are crucial to establishing the three germ layers as well as the body axes in a highly co-ordinated process. An understanding of the precise molecular and cellular mechanisms that govern these processes is a fundamental area of study in developmental biology.

The study of the early embryo was for decades reliant on *ex vivo* embryo culture. A large number of studies using various embryo isolation and culture protocols were crucial to gain a better understanding of the molecular events that govern the shaping of the embryo (Keller, 2013; Bedzhov et al., 2014; Bedzhov and Zernicka-Goetz, 2014; Schrode et al., 2014). However, the use of requirement of embryos themselves is rate limiting due to the ethical and technical challenges that accompany their usage. Recently, however, there has been a significant amount of research into the dynamic self-organising ability of cultured stem cells to recapitulate embryonic development *in vitro* (Turner et al., 2016; Simunovic and Brivanlou, 2017; Shahbazi and Zernicka-Goetz, 2018; Taniguchi et al., 2019; Zylicz, 2020; van den Brink and van Oudenaarden, 2021; Veenvliet et al., 2021; Arias et al., 2022; Amel et al., 2023). These so-called ‘stembryos’ are able to model the events that occur during pre-and peri-implantation development in a remarkably organised and consistent manner. ‘Stembryos’ have thus presented themselves as a powerful tool for investigating the mechanisms underlying early mammalian development, as they allow for unrestricted access and manipulation of developmental events and environmental cues and the tracking of cell fate dynamics without the complications of requiring live embryos.

Rapid progress has thus been made in the ‘stembryo’ field in a very short space of time, with different culture techniques able to resemble the blastocyst (Rivron et al., 2018; Sozen et al., 2019; Yu et al., 2021; Heidari Khoei et al., 2023), initiate pro-amniotic cavity formation and gastrulation (Harrison et al., 2017; Sozen et al., 2018), show signs of show signs of anterior-posterior axis elongation (van den Brink et al., 2014; Turner et al., 2017; Beccari et al., 2018), form neural tube-like structures (Veenvliet et al., 2020; Libby et al., 2021; Bérenger-Currias et al., 2022), recapitulate somitogenesis (van den Brink et al., 2020; Veenvliet et al., 2020) and even recapitulate the early stages of cardiogenesis (Rossi et al., 2021; Olmsted and Paluh, 2022). However, with the rapid advancement in this field, the complex nature of the reagents used in these protocols (which are largely shared between the various studies) has left the precise roles of these factors in germ layer differentiation and stem cell self-organisation unclear and requiring further investigation. Reagents such as N2 and B27, both initially designed to aid in the culture of various neuronal cell types and CHIR99021, a Wnt pathway activator commonly known as Chiron, are readily used in a variety of ‘stembryo’ protocols, but their individual roles in driving stem cell lineage commitment as they do in these techniques has not been fully determined. In the present study, we investigate the morphological and gene expression changes induced in embryoid bodies (EBs), the simplest three-dimensional aggregates of stem cells used to study their differentiation, exposed to commonly used ‘stembryo’ supplements to help dissect their roles in this emerging field.

## Results

### The effect of Chiron on the morphology of mES cell aggregates

The Wnt agonist Chiron is a commonly used reagent in various ‘stembryo’ culture protocols and is often included in culture via a brief ‘pulse’ together with several other supplements to drive mES differentiation. To examine the role of Chiron in mES cell differentiation we treated aggregates of ES cells with Chiron in the absence of other signalling molecules. Embryoid bodies (EBs) were subjected to a pulse of Chiron at 48 h for 24 h and were compared to EBs without Wnt agonist treatment (Fig. 1A). In the absence of Chiron, the aggregates, regardless of size, remained mostly spherical (Fig. 1B). However, the inclusion of a Chiron pulse resulted in a significant morphological change, as the aggregates elongated into a shape resembling the gastrulating embryo (Fig. 1C). To quantify the shape changes over time, the longest axis of the aggregate (the major axis) was measured, as well as the distance perpendicular to the midpoint of the longest axis (the minor axis) to determine an aspect ratio (Fig. 1A). The aspect ratio of the EBs without a Chiron did not change significantly over the 120 h period, indicating that they retained their spherical shape (Fig. 1D). EBs treated with a Chiron pulse showed a rapid increase in the aspect ratio from 48 h to 96 h, with a significantly more elongated shape compared to those without a Chiron pulse (Fig. 1D), p = 0.0021). Additionally, we observed that the shape of the aggregates varied significantly after the Chiron pulse. Aggregates that were noted to be very large as well as very small aggregates were not able to elongate even when exposed to the Wnt agonist, suggesting that the elongation process depends on the size of the aggregates. On the other hand, without a Chiron pulse, the aggregates retained their spherical shape regardless of their initial size.

**Fig. 1:**
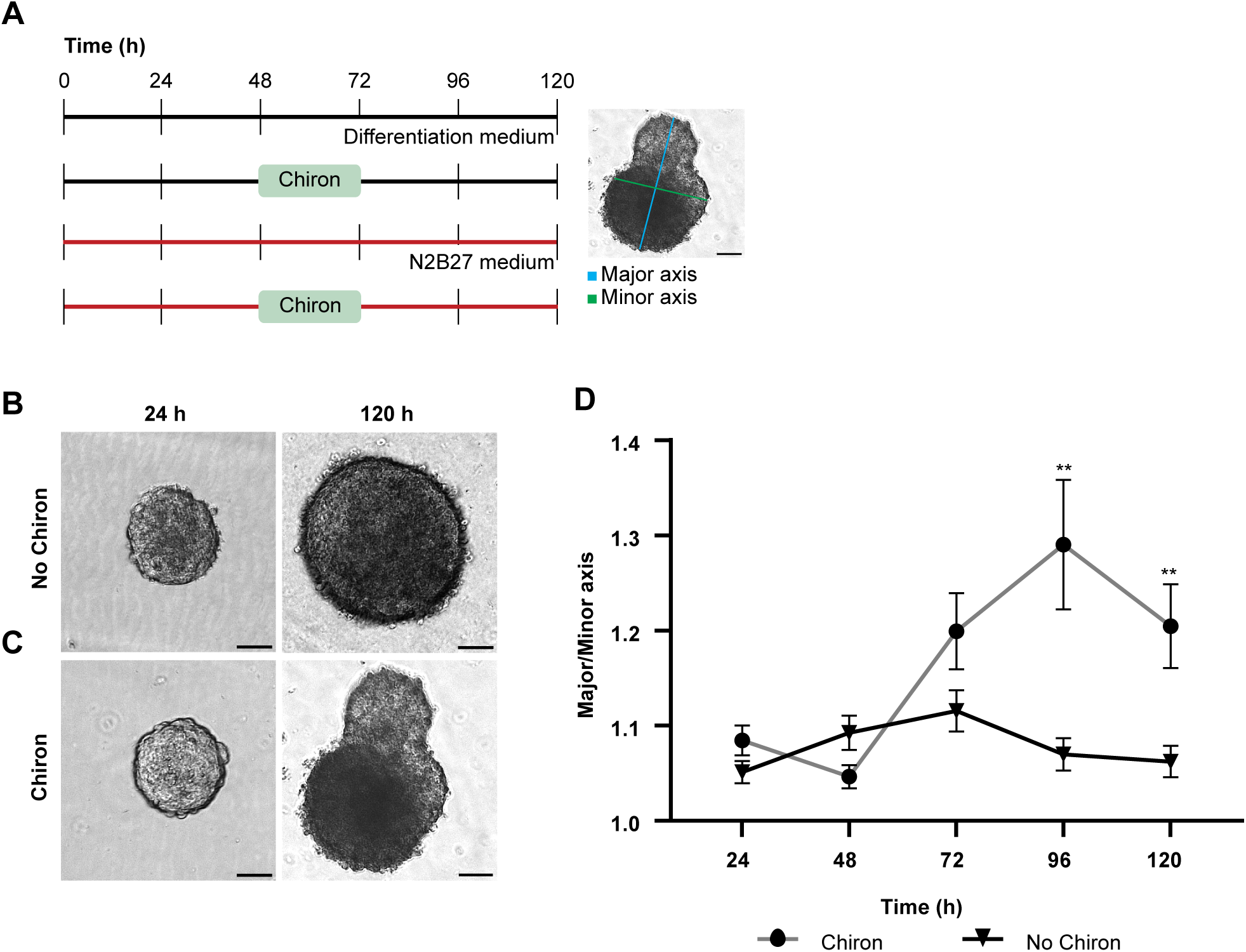
Comparative analysis of the effect of Chiron on ES cell aggregate morphology. (A) ES cell aggregates were cultured in several different conditions. In a standard differentiation medium with or without a 24 h Chiron pulse, or in N2 and B27 containing medium with or without a 24 h Chiron pulse. Elongation of aggregates was calculated by measuring the longest axis of the aggregate (the major axis; blue line) and the axis perpendicular to this measurement at its midpoint (minor axis; green line). (B) Morphology of aggregates grown in differentiation medium only. Aggregates retained their spherical shape for the duration of 120 h. (C) Morphology of aggregates grown in differentiation medium that received a Chiron pulse. Aggregates elongated following the Chiron pulse. (D)Aspect ratio of aggregates with (grey line, with dots) or without (black line with triangles) a Chiron pulse for n = 20 aggregates per condition. Error bars indicate mean ± s.e.m. For every time point, an unpaired, nonparametric Mann-Whitney test was performed and ** = p≤0.0021. Scale bar: 25 µm

### A Chiron pulse induces gastrulation and favours mesoderm and endoderm formation

The Wnt pathway plays an essential role in the induction of the primitive streak which breaks the bilateral symmetry of the embryo as epiblast cells undergo an epithelial-to-mesenchymal transition (EMT) and move inward, forming mesodermal and endodermal lineages and marking the posterior of the embryo(Tam and Loebel, 2007; Bardot and Hadjantonakis, 2020). In this study, we sought to determine the degree to which similar events could be induced in EBs via a Wnt agonist pulse. To investigate this, we measured the expression of EMT and primitive streak markers using qPCR in EBs with or without the addition of a Chiron pulse. A pulse of Chiron was sufficient to induce a down-regulation of posterior epiblast markers *Wnt3* and *ß-Catenin* expression with up-regulation of *Brachyury* and *Snai1* (Fig. 2) indicating the initiation of gastrulation and the EMT process. The up-regulation of the mesenchymal marker *Snai1* resulted in a subsequent down-regulation of epithelial marker *E-cadherin,* an epithelial marker, and is indicative of an EMT taking place (Fig. 2). Interestingly, no significant difference was noted in *Nodal* expression following a Chiron pulse (Fig. 2). Overall, these results indicate that Chiron is able to drive an EMT in ES cells in culture.

**Fig.2:**
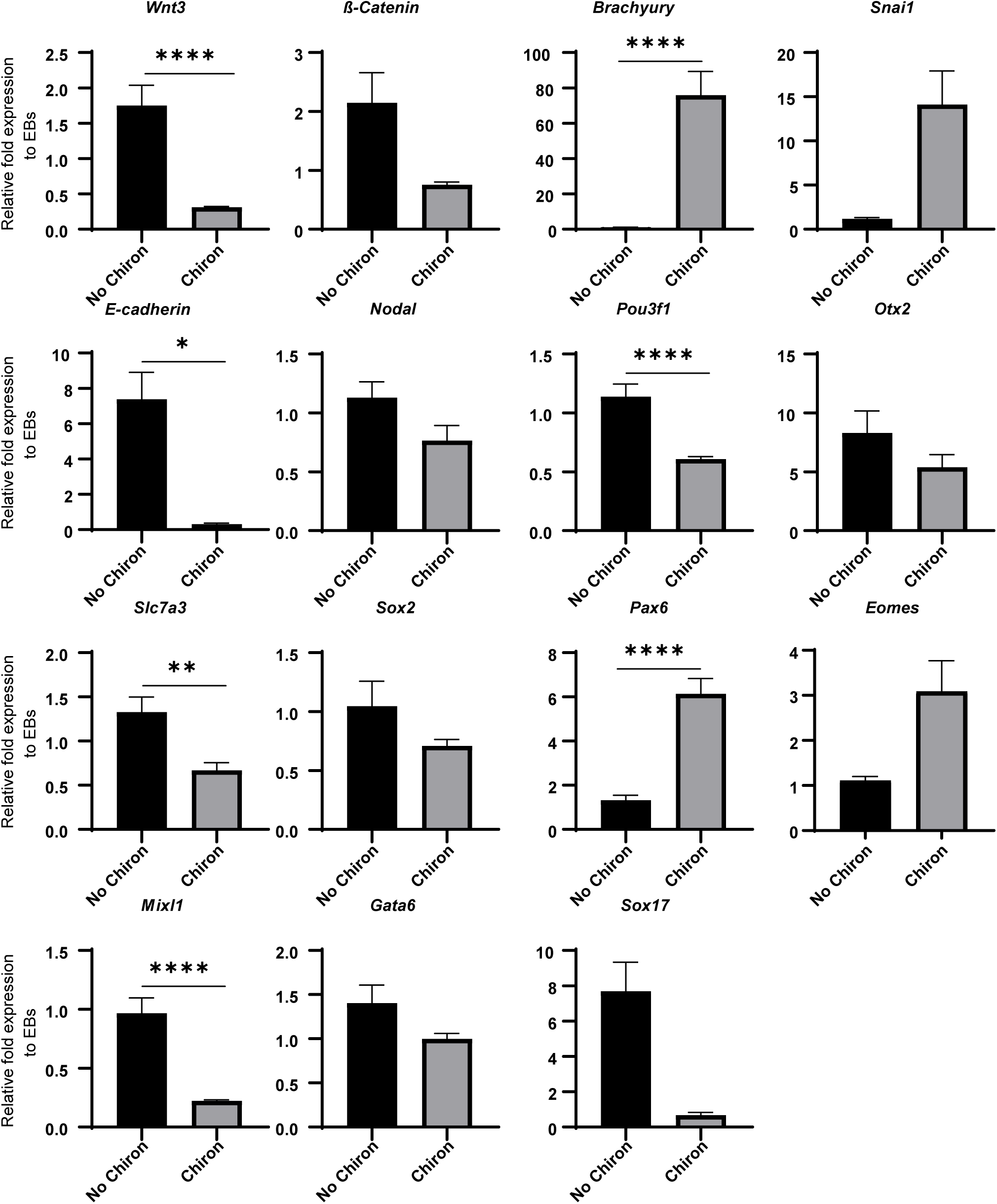
Effects of a Chiron pulse on gene expression of aggregates grown in differentiation medium. QPCR was performed on reverse transcribed RNA extracted at 120 h for n=2 biological replicates. Primitive streak markers: *Wnt3, ß-catenin, Brachyury* and *Nodal.* EMT markers: *E-cadherin* and *Snai1*. Anterior/neurectoderm markers: *Pou3f1, Otx2, Slc7a3, Sox2* and *Pax 6*. Mesendoderm markers: *Eomes* and *Mixl1.* Endoderm markers: *Gata6* and *Sox17*. Relative fold expression to EBs without Chiron. Error bars indicate mean ± s.e.m. An unpaired, nonparametric Mann-Whitney test was performed and * = p≤0.0332, ** = p≤0.0021, *** = p≤0.0002, **** = p< 0.0001.

Primitive streak formation is a crucial event in establishing the embryonic germ layers. To assess the degree to which a Wnt agonist can influence lineage commitment, the expression of various germ layer markers was quantified using qPCR with and without a Chiron pulse. Expression levels of anterior/neuroectoderm markers were generally significantly down-regulated following a pulse of Chiron, with decreased expression noted in *Pou3f1, Otx2, Slc7a3* and *Sox2* (Fig. 2). This is consistent with the known posteriorising effects of Wnt signalling on the embryo. However, *Pax6,* a key transcription factor in central nervous system development, was significantly up-regulated following the induction of Wnt signalling (Fig. 2, p<0.0001). The mesendoderm marker *Eomes* showed a significant up-regulation following a Chiron pulse (Fig. 2). However, another mesendoderm marker *Mixl1* was significantly down-regulated following a Wnt pulse (Fig. 2). The endoderm markers *Gata6* and *Sox17* were also down-regulated, although not significantly (Fig. 2). Overall, these results show that a Chiron pulse is sufficient to drive mesoderm and endoderm differentiation at the expense of ectoderm formation in mES cells.

### N2B27 contrasts posteriorising signals of Chiron

N2 and B27 are critical components of various ‘stembryo’ culture protocols (van den Brink et al., 2014; Harrison et al., 2017; Turner et al., 2017; Beccari et al., 2018), but the precise role of these complex supplements, particularly with the addition of a pulse of Wnt agonist, requires further investigation. To explore the effect of N2B27 on the expression of EMT, primitive streak formation, and germ layer induction, we investigated the gene expression of various relevant markers using qPCR in EBs cultured in N2B27 medium with and without a Chiron pulse.

The addition of a Chiron pulse had a different effect on *Wnt3* expression in the presence and absence of N2 and B27. In the presence of N2B27 medium, *Wnt3* expression was significantly up-regulated by the Chiron pulse (Fig. 3), while it was down-regulated without N2B27 medium (Fig. 2). While a down-regulation of *ß-catenin* and an up-regulation of *Brachyury* was observed following a Chiron pulse in N2B27 medium (Fig. 3) similar to what was seen before (Fig. 2), the fold expression was significantly reduced in N2B27 containing medium. Both *Nodal*, a primitive streak and mesoderm marker, and *E-cadherin*, an epithelial marker, were significantly up-regulated following a Chiron pulse in N2B27 medium (Fig. 3), again contrasting the expression changes seen without N2 and B27 (Fig. 2). There was a significant up-regulation of the EMT marker, *Snai1,* following a pulse of the Wnt agonist in N2B27 medium (Fig. 3) coinciding with the effect of Chiron on *Snai1* seen before (Fig. 2).

**Fig. 3:**
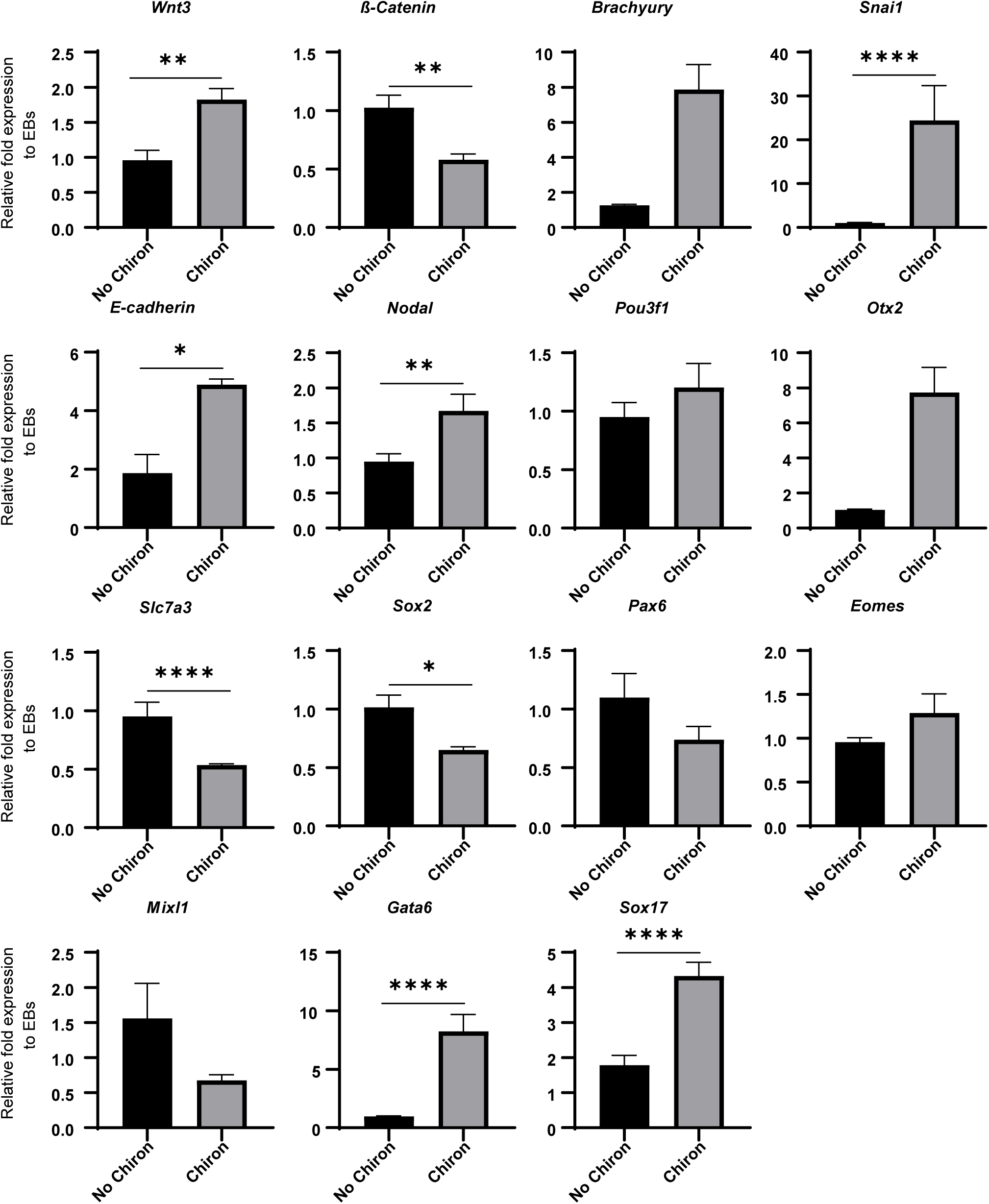
Effects of a Chiron pulse on gene expression of aggregates cultured in N2B27 medium. QPCR was performed on reverse transcribed RNA extracted at 120 h for n = 2 biological replicates. Primitive streak markers: *Wnt3, ß-catenin, Brachyury* and *Nodal.* EMT markers: *E-cadherin* and *Snai1.* Anterior/neurectoderm markers: *Pou3f1, Otx2, Slc7a3, Sox2* and *Pax6.* Mesendoderm markers: Eomes and Mixl1. Endoderm markers: *Gata6* and *Sox17*. Relative fold expression normalised to EBs without Chiron. Error bars indicate mean ± s.e.m. An unpaired, nonparametric Mann-Whitney test was performed and * = p≤0.0332, ** = p≤0.0021, *** = p≤0.0002, **** = p< 0.0001.

There was a significant up-regulation of the early anterior marker *Otx2* (Fig. 3) and a down-regulation of both *Slc7a3* and *Sox2*, markers of the anterior epiblast and neural stem cells, respectively, following a Chiron pulse in N2B27 medium. There were no significant effects on *Pou3f1* and *Pax6* expression following a Chiron pulse in N2B27 medium (Fig. 3). Both mesendoderm markers, *Eomes* and *Mixl1*, did not have any significant differences in expression in N2B27 medium following a Chiron pulse (Fig. 3) although it was noted that the trend of both these markers was similar to what was seen in the absence of N2 and B27 following Wnt signalling (Fig. 2). Interestingly, the expression of both endoderm markers investigated, *Gata6* and *Sox17,* was significantly up-regulated following Wnt induction in N2B27 medium (Fig. 3). Together these results suggest a complex interplay between N2, B27 and Chiron in driving gene expression changes in ES cell aggregates in culture.

### BMP4 has contrasting effects to Wnt signalling on the elongation of ES cell aggregates in culture

BMP4 is crucial for initiating anterior-posterior patterning and gastrulation in the mouse embryo (Arnold and Robertson, 2009b; Bardot and Hadjantonakis, 2020). Recently, it has been shown that exposure to a BMP4 signalling centre is sufficient to drive embryonic morphogenesis in culture (Xu et al., 2021). To examine the role of BMP4 signalling on mES cell differentiation we treated ES cell aggregates with pulses of both BMP4 and Chiron (BMP4 + Chiron) or combined ES aggregates with a BMP4 signalling centre (BMP4 SC) (Fig. 4A). We compared the aggregates with EBs treated with just a Chiron pulse as above with all culture taking place in N2B27 containing medium (Fig. 4A).

**Fig. 4:**
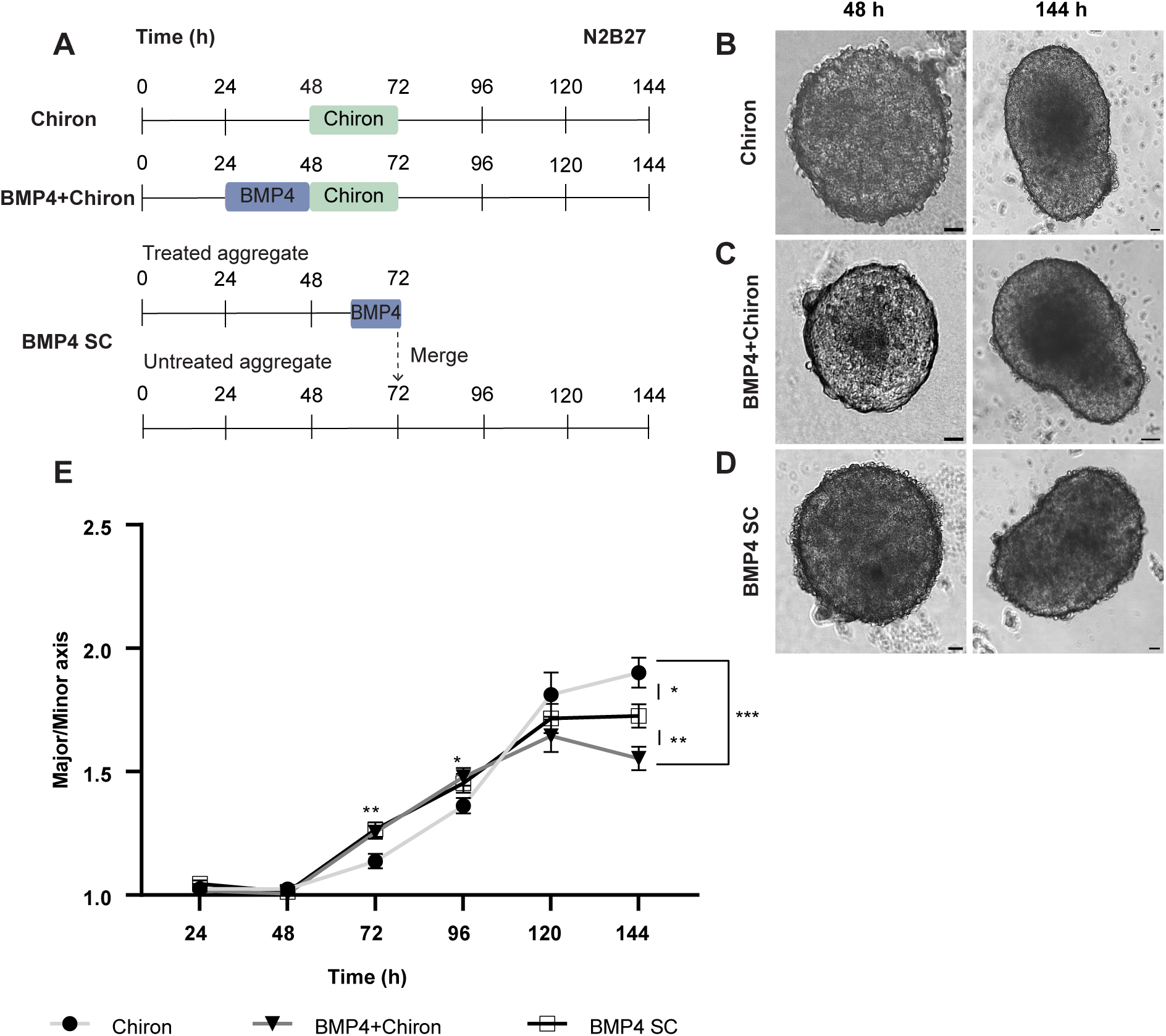
Comparative analysis of the effect of BMP4 on the ES cell aggregate morphology. (A) ES cell aggregates were cultured in several different conditions. Chiron aggregates were cultured in N2B27 medium and received a 24 h Chiron pulse on the third day of culture. BMP4 + Chiron aggregates received a 24 h BMP4 pulse followed by a 24 h Chiron pulse. BMP SC aggregates were made by merging a smaller aggregate, which had been treated with BMP4 for 8 h, with a larger, untreated aggregate on the fourth day of culture. (B) Morphology of Chiron aggregates at 48 h and 144 h. Aggregates elongated following the Chiron pulse. (C) Morphology of BMP4 + Chiron aggregates at 48 h and 144 h. Aggregates were able to elongate effectively but not as much as Chiron aggregates. (D) Morphology of BMP4 SC aggregates at 48 and 144 h. Aggregates were able to elongate but retained an oval shape. (E) Aspect ratio of Chiron (light-grey line, with dots), BMP4 + Chiron (grey line with triangles) and BMP SC (black line with stars) aggregates for n = 15 aggregates per condition. Error bars indicate mean ± s.e.m. A one-way ANOVA with Dunnett’s Multiple Comparison’s post-test was carried out and * = p≤0.0332, ** = p≤0.0021, *** = p≤0.0002, **** = p< 0.0001. Scale bar: 25µm

At 48 h, all aggregates were roughly spherical in shape and had doubled from initial seeding size (Fig. 4B-D). EBs exposed to Chiron began to elongate by 72 h and continued to do so throughout the culture period (Fig. 4B and E). While EBs treated with both pulses of both BMP4 and Chiron did break symmetry and elongate (Fig. 4C), by 144 h there was a significant reduction in aspect ratio compared to those treated with Chiron alone (Fig. 4E). Similarly, EBs treated with a BMP4 signalling centre were also found to be more oval in shape than those treated with a Chiron pulse (Fig. 4D) and had an even more significantly reduced aspect ratio by 144 h (Fig. 4E). Interestingly, while by 144 h the Chiron aggregates had, as mentioned, elongated more than the other conditions, the aspect ratio of BMP4 aggregates (both those receiving a pulse or a signalling centre) was significantly higher at 72h and 96 h when compared to the Chiron aggregates (Fig. 4E). These results suggest that BMP4 and Wnt signalling have contrasting roles to play in the rate and degree to which ES cell aggregates can elongate.

### Localised BMP4 signalling allows the expression of anterior neural markers in cell aggregates

As the inclusion of BMP4 had a clear effect on ES aggregate morphology we next investigated the effects of our culture conditions on the expression of various germ layer markers, using qPCR. Exogenous BMP4 significantly increased *Bmp4* mRNA expression under BMP4+Chiron conditions as expected (Fig. 5A). Interestingly, BMP4 SC aggregates had similar levels of *Bmp4* mRNA as the Chiron treated aggregates, despite the latter not receiving any exogenous BMP4 (Fig. 5A). Both conditions exposed to a Wnt agonist showed an increase in *Wnt3* mRNA expression as expected, while BMP4 SC aggregates did not have significant levels of *Wnt3* compared to the control (Fig. 5A). Interestingly, conditions that included BMP4 had significantly up-regulated levels of *E-cadherin* compared to Chiron-treated EBs with the BMP4 SC conditions expressing the highest levels of *E-cadherin* (Fig. 5A). *Brachyury* and *Mixl1* expression were expressed in all conditions without high levels of variation, while there was a striking effect on *Sox17* expression with a significant up-regulation in cultures containing BMP4.

**Fig. 5:**
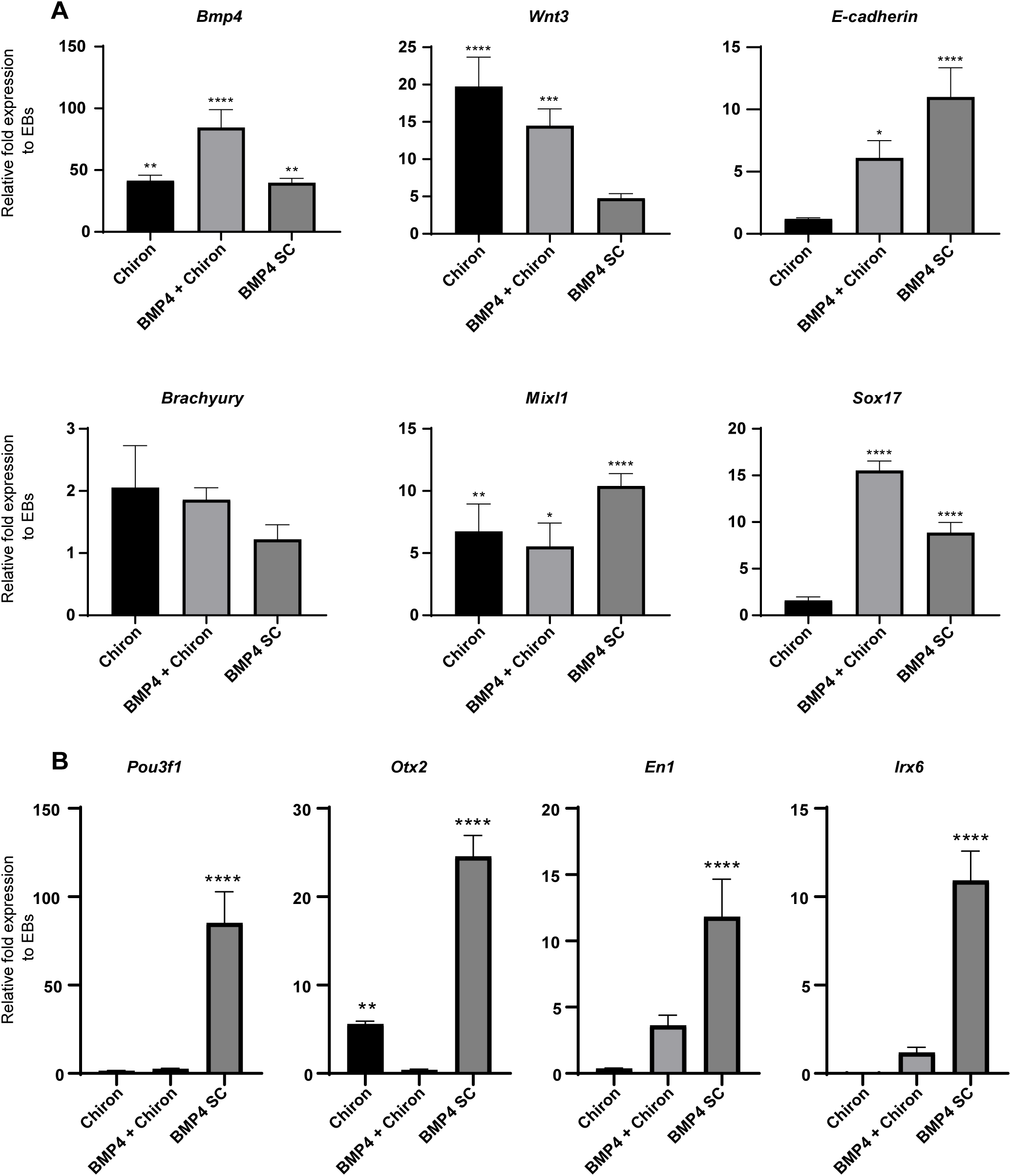
Effects of BMP4 on gene expression of aggregates. (A) QPCR was performed on reverse transcribed RNA extracted at 144 h for n=2 biological replicates. posterior and primitive streak markers: *BMP4* and *Wnt3*. EMT marker: *E-cadherin*, mesoderm markers: *Brachyury* and *Mixl1* and endoderm marker: *Sox17*. (B) Anterior neuroectodermal markers were highly expressed in BMP4 SC aggregates. Relative fold expression normalised to EBs without Chiron. Error bars indicate mean ± s.e.m. A one-way ANOVA with Dunnett’s Multiple Comparison’s post-test was carried out and * = p≤0.0332, ** = p≤0.0021, *** = p≤0.0002, **** = p< 0.0001.

Most strikingly, all the anterior neural markers investigated, *Pou3f1, Otx2, En1, and Irx6,* were all highly expressed in BMP SC cultures with this same effect not being observed under conditions with a BMP4 pulse (Fig. 5B).

## Discussion

The ability of stem cells to self-organise in culture and form complex, embryo-like structures presents a non-invasive, scalable, and ethically viable way to investigate gastrulation and germ layer formation in mammals. Many ‘stembryo’ culture systems make use of N2 and B27 (van den Brink et al., 2014; Baillie-Johnson et al., 2015; Turner et al., 2017; Beccari et al., 2018), supplements originally designed to specifically support neuronal cell culture (Bottenstein and Sato, 1979; Brewer et al., 1993). B27, which contains retinyl acetate, is particularly important for the formation of neuronal tissue in the posterior hindbrain and the anterior region of the spinal cord (Brewer et al., 1993; Ross et al., 2000), while N2 contains insulin, selenium and transferrin and has been shown to support neurogenesis (Bottenstein and Sato, 1979).When mES cells are cultured in medium containing both N2 and B27 they differentiate towards a neural fate (Ying et al., 2003; Abranches et al., 2013). ‘Stembryo’ culture protocols also make liberal use of a potent Wnt agonist CHI99021, commonly known as Chiron, in a pulsatile manner (van den Brink et al., 2014; Baillie-Johnson et al., 2015; Turner et al., 2017; Beccari et al., 2018). Wnt signalling is known to be involved in initiating gastrulation through up-regulation of the primitive streak marker Brachyury in the embryo (Dunty et al., 2008; Wang et al., 2012; Arkell et al., 2013) and has a similar effect in stembryo culture (van den Brink et al., 2014; Baillie-Johnson et al., 2015; Turner et al., 2017; Beccari et al., 2018). BMP4 is also critical for the initiation of the primitive streak. BMP4 from the extraembryonic ectoderm up-regulates the expression of Wnt3 and Nodal in the posterior region of the epiblast, which initiates this symmetry breaking process and creates a feed-forward signalling complex (Perea-Gomez et al., 2004; Rivera-Perez and Magnuson, 2005; Arnold and Robertson, 2009a). The expression of these transcription factors is restricted to the posterior region by antagonists expressed in the anterior epiblast (Thomas and Beddington, 1996; Kumar et al., 2015; Antonica et al., 2019). Under the influence of anterior transcription factors, this region will form the neural plate, giving rise to the brain. Treating stem cell aggregates with BMP4 has been shown to induce the expression of Wnt and Nodal (Xu et al., 2021). When these BMP4 aggregates are merged with a second, larger, untreated aggregate, it is able to act as a signalling centre to initiate gastrulation in the now merged ‘stembyro’ (Xu et al., 2021). The combination of these various signalling pathways, induced in stem cells using a variety of supplements and ligands, has resulted in the rapid development of various ‘stembryo’ culture systems. This novel field continues to deliver critical insights into the earliest organisational events that occur during mammalian development in a short period of time. However, the rapid development of this field has resulted in the precise effects of some of these various culture supplements on stem cell differentiation remaining undefined. Here, we treated embryoid bodies with various ‘stembryo’ culture reagents to begin investigating their precise effects on stem cell aggregate morphology and gene expression.

Our results show that Chiron makes stem cell aggregate elongation a more robust process, an observation supported by other studies (van den Brink et al., 2014; Turner et al., 2017). Interestingly, BMP4 hindered the elongation of stem cell aggregates, even in the presence of Chiron. This is consistent with previous studies that found that BMP4 can inhibit axial elongation in the mouse embryo (Turner et al., 2017; Girgin et al., 2021).

We found that the addition of Chiron resulted in a decrease in *Wnt3* and *ß-catenin* mRNA in EBs. As Chiron is known to act downstream of the Wnt/ß-catenin pathway, we hypothesise that the addition of Chiron in culture initiates a negative feedback loop in this pathway, down-regulating the expression of *Wnt3* and *ß-catenin*. We expected that BMP4 together with Chiron would enhance the expression of *Wnt3* mRNA by recapitulating the feed-forward loop normally found in the posterior region of the embryo (Liu et al., 1999; Tam and Loebel, 2007; Arnold and Robertson, 2009b). Although EBs treated with both BMP4 and Chiron had significantly higher levels of *Wnt3* mRNA compared to untreated EBs, *Wnt3* expression was reduced compared to aggregates treated with just Chiron, indicating that BMP4 had hindered *Wnt3* expression in differentiation cultures. Chiron aggregates were also found to have high levels of *Bmp4* compared to untreated aggregates even in the absence of exogenous BMP4 treatment. It may be that Wnt3 activation, through a Chiron pulse, resulted in an increase in *Bmp4* expression, mimicking the BMP4/Nodal/Wnt signalling complex that occurs in the posterior proximal region of the embryo during primitive streak formation and gastrulation (Liu et al., 1999; Tam and Loebel, 2007; Arnold and Robertson, 2009b). Similarly, it can be argued that the signalling centre treated with BMP4 under BMP SC conditions, which has been shown to have an induced expression of Nodal and Wnt3 (Xu et al., 2021), interacts with the untreated aggregate and induces the expression of *Bmp4*.

In our study, the addition of Chiron was able to down-regulate *E-cadherin* expression and up-regulate *Snai1* expression as occurs in the embryo during EMT (Kim et al., 2017), thereby suggesting that Chiron facilitates this process in mES cell aggregates. Chiron acted as a posteriorising signal directing differentiation towards the mesendoderm lineage, as observed with decreased expression of anterior markers and increased expression of the primitive streak marker and mesendoderm markers *Brachyury* and *Eomes*. We also observed a decrease in the endoderm markers *Sox17* and *Gata6* after the EBs were treated with a Chiron pulse, which may suggest that Chiron directs mesendoderm progenitors towards the mesoderm rather than the endoderm lineage (Ying et al., 2008), although we should expect to see endoderm progenitors in our structures (van den Brink et al., 2014; Turner et al., 2017). We found that aggregates exposed to either a BMP4 pulse or signalling had a significant increase in the expression of the definitive endoderm marker Sox17, and this is supported by previous work that has shown that BMP4 functions to specify the definitive endoderm lineage (Teo et al., 2011; Teo et al., 2014). The definitive endoderm forms an epithelial layer that covers the epiblast and is associated with high levels of E-cadherin (Nowotschin et al., 2019). Studies have shown that stem cell aggregates form an endoderm-like layer on their surface and these cells co-express Sox17 and E-cadherin, resembling the definitive endoderm forming on the outside of the mouse embryo (van den Brink et al., 2014; Hashmi et al., 2022). It follows that BMP treatment of EBs is driving differentiation towards the endoderm lineage.

Interestingly, *Pax6*, a gene involved in the formation of the central nervous system (Osumi et al., 2008) was up-regulated in our Chiron-treated aggregates. Various studies have shown that Wnt signalling upregulates *Pax6* expression in the mouse brain and its outgrowths, including the cerebellum (Yeung et al., 2016), the optic cup(Canto-Soler and Adler, 2006), and the olfactory epithelium (Collinson et al., 2003). Although our aggregates have not reached these advanced stages of development, increased expression of *Pax6* may be an indicator of precursor cells that would eventually develop into these features.

Wnt signalling plays a broad role in the developing mouse embryo and regulates different downstream effectors depending on the tissue and stage of development. In our study, we found that Chiron acts downstream of the Wnt/ß-catenin pathway and serves to make axial elongation a more reproducible event in mES cell aggregates. As a result of Chiron signalling, Brachyury expression is up-regulated, and the primitive streak is initiated. Differentiation is primarily directed toward the mesoderm lineage as a result of activated Wnt signalling, however, due to the complex role of the Wnt signalling pathway in differentiation, there may be an increase in expression of non-mesodermal markers, such as Pax6. We also hypothesise that Chiron, as a downstream effector of the Wnt pathway, may regulate the expression of Wnt proteins through a negative feedback loop. We found that BMP4, which, in the embryo, acts to enhance the effects of Wnt signalling, appears to hinder the effect of Chiron by preventing the elongation of aggregates that have been subjected to a Chiron pulse. In contrast to the mesoderm-inducing effects of Chiron, BMP4 favours endoderm differentiation. Mesoderm and endoderm cells share a common precursor: mesendoderm cells. In the embryo, the path of differentiation is dictated by the concentration of various transcription factors, including BMP4 and Nodal (Vincent et al., 2003; Bardot and Hadjantonakis, 2020). To faithfully recreate these events *in vitro*, optimisation of the dosage and timing of exogenous factors such as Chiron and BMP4 is needed in order to recapitulate the body plan in mES cells.

Cell aggregates exposed to high Wnt signalling, in the form of a Chiron pulse, lack brain and head structures, and resemble the post-occipital region of embryos (van den Brink et al., 2014; Turner et al., 2017; Beccari et al., 2018). We found that the anterior markers *Pou3f1*, *Otx2* and *Slc7a3* were down-regulated with the addition of a Chiron pulse in our EBs, thereby suggesting that Chiron hindered anterior neural fate commitment. On the other hand, it has previously been shown that when mES cell monolayers are cultured in N2B27 medium they differentiate into glial subtypes such as astrocytes and oligodendrocytes (Ying et al., 2003; Abranches et al., 2009). Therefore, we cultured EBs in N2B27 medium with and without a Chiron pulse to investigate if this would balance anterior vs posterior differentiation in our stem cell aggregates and recreate a more complete body plan with anterior and posterior structures. We noted an increase in *Wnt3* expression after a Chiron pulse in N2B27 medium, in contrast to the effect of Chiron on *Wnt3* without the presence of N2 or B27. We hypothesise that this reversal of effect is due to the complex nature of the cell culture medium, with various components, such as retinyl acetate in B27, that can modulate the response of cells to Chiron and result in an increase in *Wnt3* expression. We hypothesised that since Chiron acts downstream of the Wnt/ß-catenin pathway, it is able to up-regulate the expression of downstream targets of this pathway, which, in turn, work in a negative feedback loop to down-regulate the expression of *Wnt3*. Studies have shown that retinoic acid (a retinyl acetate metabolite) inhibits downstream Wnt/ß-catenin events (Hu et al., 2013; Osei-Sarfo and Gudas, 2014) but increases the expression of canonical Wnt ligands: Wnt2, Wnt3a and Wnt8a (Osei-Sarfo and Gudas, 2014). Therefore, it may be possible that the retinyl acetate in our medium reduces the expression of downstream targets of the Wnt pathway which would affect the regulatory loop and result in an up-regulation of the Wnt3 gene. *Brachyury* remained up-regulated in the presence of N2B27 and Chiron, although not as significantly as expression in the absence of N2B27, suggesting that N2B27 may balance the expression of mesoderm or posterior markers with anteriorising signals (Thomson et al., 2011; Turner et al., 2014b) possibly from the effects of retinyl that is essential for neocortex formation (Haushalter et al., 2017). We observed an increase in the expression of the anterior epiblast markers *Pou3f1* and *Otx2* in the presence of N2B27 and Chiron, contradicting the assumption that Chiron increases the expression of posterior markers and decreases the expression of anterior markers (Turner et al., 2014a). It may be that N2 and B27 reverse the posteriorising effects of Chiron by enhancing the expression of anterior markers. The results suggest that the addition of Chiron to stem cell aggregates promotes posterior neural fate commitment and inhibits the development of brain and head structures. However, the use of N2B27 medium appears to balance the expression of posterior markers with anteriorizing signals, resulting in the expression of anterior markers. Therefore, tightly regulating the timing and specific anteriorising and posteriorising signals in ‘stembyro’ culture systems will allow the recreation of a more complex *in vitro* embryo model.

We found that introducing BMP4 as a spatially restricted signalling centre in aggregates resulted in significant up-regulation of neural markers, including *Pou3f1, En1, Irx6* and *Otx2*. There was a notable and distinct increase in all of these markers in the BMP4 signalling centre aggregates compared to those that just received a BMP4 pulse. This is significant evidence that it is not only the concentration at which supplements are added to cell culture that is critical, but the way they are introduced that can direct stem cell differentiation in ‘stembryo’ models. The increase in the expression of anterior neural markers under BMP4 SC conditions confirms the ability of this model to recreate the anterior embryonic axis. In this model, the signalling centre is exposed to BMP4, which induces the expression of Nodal and Wnt3. Once this signalling centre merges with the untreated aggregate, it serves as the posterior pole of the aggregate with high concentrations of Nodal and Wnt3, as occurs in the embryo. A morphogen gradient is created, and the region farthest away from the signalling centre will receive the least Nodal and Wnt3 activity and will serve as the anterior pole. In the embryo, the anterior visceral endoderm is responsible for secreting the Nodal and Wnt3 agonists, Dkk1, Cer1, and Lefty, which restrict the activity of Nodal and Wnt to the posterior region (Arnold and Robertson, 2009b; Arkell and Tam, 2012). This model does not contain the anterior visceral endoderm and is based solely on the engineered signalling gradient and serves to demonstrate the importance of a morphogen gradient in creating the body’s axes. However, we cannot ignore the roles of Dkk1, Cer1, and Lefty1 in development, and future studies should attempt to create a morphogen gradient of these anterior markers in stem cell-based models and combine anterior and posterior signalling molecules in one system.

A notable limitation of our study is the prevalence of FBS in our culture medium. Besides the fact that FBS contains undefined growth factors and may be subject to batch-to-batch variability, it has also been shown that EBs cultured in serum-containing medium have been shown to have a localised expression of Wnt and Nodal, which in turn results in increased Brachyury (Ten Berge et al., 2008). FBS also contains BMP4, which may contribute to the induction of Brachyury expression and the initiation of primitive streak formation and gastrulation in the absence of Chiron. Future ‘stembryo’ studies should focus on excluding undefined components and replacing serum entirely to improve experimental reproducibility. While beyond the scope of this study, future work will test the effects of N2 and B27 individually on stem cell morphology and gene expression especially on how this will affect anterior/neuroectoderm markers. N2 and B27 are consistently used together in ‘stembryo’ cultures, and it would be interesting to note their effects when segregated.

Our work described here provides an examination of how common ‘stembryo’ culture reagents, Chiron, N2, B27, and BMP4 signalling impact the morphology and gene expression of stem cells in the simplest 3D differentiation model. Specifically, we found that Chiron treatment of mES cells makes EMT and gastrulation more reproducible, that the Wnt pathway plays a critical role in the posterior region of the embryo during primitive streak formation, and that N2B27 medium can counterbalance the posteriorising effects of the Wnt pathway and promote lineage commitment to the neuroectoderm. Critically, we find that a group of localised cells exposed to BMP4 can effectively drive neural differentiation in stem cell aggregates. Finally, the use of a signalling centre to establish a morphogen gradient can lead to the more prominent establishment of anterior and posterior identity, resulting in a more accurate representation of the developing body and could be a key technique in the ‘stembryo’ field moving forward.

## Materials and methods

### mES Cell culture

A feeder layer of inactivated murine embryonic fibroblast cells was cultured in 6-well plates on gelatin (0.1%, Sigma-Aldrich G7041-100G) at a seeding density of 1.6 x 10^4^ cells/cm^2^ in base medium consisting of DMEM (1x) +GlutaMAX™ (Gibco 10566016), 15% FBS (Gibco 10493106), 0.05 mM β-mercaptoethanol (Gibco 21985023), 1% PenStrep (100 U penicillin/ 0.1 mg/ml streptomycin, Gibco 15140122). Two h prior to seeding mES cells, the medium was changed to the same base medium with the addition of CHIR99021 (Chiron, 3 µM, Sigma-Aldrich SML1046-5MG), PD0325901 (1 μM, Sigma-Aldrich PZ0162-5MG) and Leukaemia inhibitory factor (10 ng/ml, LIF, Thermo Fisher Scientific A35934). 129/Ola mES cells were seeded on the feeder layer at a density of 7 x 10^3^ cells/cm^2^. Medium was changed daily, and cells passaged every other day.

### mES Differentiation

mES colonies were treated with Dispase II (5mg/ml, Sigma-Aldrich D4693-1G) at 37°C for 20 min, followed by inactivation of Dispase and transfer of the cell suspension to a 15 ml tube. The cells were left to settle at the bottom for 10 min, after which time the supernatant was removed and the cells were resuspended in base medium, referred to as differentiation medium, or N2B27, which is made up of differentiation medium supplemented with 1% N2 supplement (Gibco 17502048), 1% B27 plus supplement. (Gibco A3582801). The cells were triturated gently to break the clusters into smaller aggregates. The contents of the tube were then transferred to a 6 cm bacteriological plate and an additional 5 ml of differentiation medium or N2B27 was added to the plate. Alternatively, to form individual aggregates, dissociated mES cells were resuspended at a concentration of 6.25×10^3^ cells/ml in N2B27 and 40 µl droplets of were pipetted into each well of a non-tissue-culture treated U-bottomed 96-well plate (Greiner Bio-One 650185). For experiments requiring a Chiron pulse, CHIR99021 was added at a final concentration of 3 µM as added 48 h to 72 h post aggregate formation. For experiments with a BMP and Chiron pulse, 24 h post individual aggregate formation, BMP4 (Gibco PHC9534) was added to each well of the U-bottomed 96-well plate at a final concentration of 1 ng/ml. After 24 h, the medium was replaced with N2B27 supplemented with Chiron at a final concentration of 3 µM. To create a BMP signalling centre, 40 μl droplets of a suspension of 3.75×10^3^ mESC cells/ml in N2B27 medium were pipetted into each well of a U-bottomed 96-well plate and left to settle for 40 h after which, BMP4 at a final concentration of 1ng/ml was introduced into each well for a total of 8 h. Simultaneously, a second aggregate of mES cells was created by placing 40 μl droplets of a 2.5×10^4^ mES cells/ml suspension in N2B27 medium pipetted into each well of a U-bottomed 96-well plate and left to settle for 48 h. Then, both these aggregates were merged 48 h after their initial formation.

### RNA extraction and cDNA synthesis

RNA was extracted at 120 h or 144 h post aggregation using the High Pure RNA Isolation Kit (Roche 11828665001) according to the manufacturer’s instructions. Due to the limited amount of material isolated from each experiment, two experiments were pooled to create each of the biological replicates. The ImProm-II™ Reverse Transcription System (Promega) was used for cDNA synthesis. Briefly, a minimum of 1 µg of RNA was combined with 1 µl of Oligo (dT)15 primer (Promega, C110B) and nuclease-free water to make up a total of 5 µl and incubated at 70°C and 4°C for 5 min each. PCR master mix was added to each sample:6.1 µl nuclease-free water, 4 µl 5x reaction buffer (Promega, M289A), 2.4 µl MgCl2 (Promega, A351H), 1 dNTP mix (Promega, C114B), 0.5 µl Recombinant RNasin® Ribonuclease Inhibitor (Promega, N251A), 1 µl Reverse Transcriptase (Promega, M314A). Thermal cycling was as follows: 25°C for 5 min, 42°C for 60 min and 70°C for 15 min.

### qPCR

Quantitative PCR was performed on the StepOnePlus™ Real-Time PCR System using SYBR green PCR Master-Mix (Thermo-Fisher Scientific 4368708). Primer sequences can be found in supplementary material Table S1. Briefly, 2 µl of cDNA (1:1 dilution with nuclease-free water) was mixed with 8 µl master which consisted of: 5 µl SYBR green Master-Mix, 0.4 µl of 10 µM forward and reverse primer mixture (5 µl of 100 µM stock with 40 µl nuclease-free water), and 2.6 µl nuclease-free water. Each reaction had three technical replicates; see supplementary material Table S2 for run parameters. *Gapdh* was used as a housekeeping gene to calculate relative expression via the 2-ΔΔCt method. Expression was normalised to EBs cultured in either differentiation medium or N2B27 medium that did not receive a Chiron pulse. MS Excel was used for data analysis, and GraphPad Prism8 was used for statistical analysis and graph generation.

### Image analysis

Images were taken using EVOS™ M5000 Imaging System microscope (Thermo Fisher Scientific, USA) at 10X magnification. All images were processed using ImageJ. For morphology measurements, the outlines of the entire aggregate were traced manually. The minor and major axial lengths were determined using the line tool. The elongation index was calculated by dividing the major axial length by the minor axial length.

### Statistical analysis

All statistical analyses were performed using the built-in functions of GraphPad Prism 8.4.2 (679). Statistical analysis of the RT-qPCR data was performed on two pooled biological replicates due to limited biological material. When appropriate, an unpaired, nonparametric Mann-Whitney test was performed. Alternatively, a one-way ANOVA with Dunnett’s Multiple Comparison’s post-test was carried out. In all cases, the error bars represent the mean ± standard error of the mean (mean ± s.e.m.). P values are represented as follows: ns = p≤0.1234, * = p≤0.0332, ** = p≤0.0021, *** = p≤0.0002, **** = p< 0.0001.

## Acknowledgements

We thank Frank Brombacher for the mES cells and Thulisa Mkatazo for assistance with generating inactivated murine embryonic fibroblasts.

## Author Contributions

M.G. conceived the project. A.A and S.R carried out the experiments. A.A., S.R., and M.G. wrote the paper.

## Data availability

All relevant data can be found within the article and its supplementary information.

